# Spatio-temporal Dynamics of the Patterning of *Arabidopsis* Flower Meristem

**DOI:** 10.1101/2020.12.02.375790

**Authors:** José Díaz, Elena R. Álvarez-Buylla

## Abstract

The qualitative model presented in this work recovers the onset of the four fields that correspond to those of each floral organ whorl of *Arabidopsis* flower, suggesting a mechanism for the generation of the positional information required for the differential expression of the A, B and C identity genes according to the ABC model for organ determination during early stages of flower development. Our model integrates a previous model for the emergence of WUS pattern in the apical meristem, and shows that this pre-pattern is a necessary but not sufficient condition for the posterior information of the four fields predicted by the ABC model. Furthermore, our model predicts that LFY diffusion along the L1 layer of cells is not a necessary condition for the patterning of the floral meristem.

## 1 Introduction

Morphogenesis occurs in plants during their whole life-cycle, with aerial and root structures forming from groups of undifferentiated or stem cells within niches found in the apical meristems in the shoot and root tips, respectively. When a plant becomes florally induced the shoot apical meristem (SAM) switches from a vegetative to an inflorescence meristem. The vegetative meristem only produces leaves as lateral organs, while the inflorescence one produces flowers that arise from its flanks in a spiral arrangement. Flowers develop from the floral meristems and in *Arabidopsis* the four sepal primordia are the first to arise from the outermost of the flower meristem (18 hrs after floral primordial formation), and the remaining floral meristem interior differentiates into the other whorls with the gynoecial primordium forming in the center of the floral primordium. At least four genes are necessary for the specification of floral meristem identity in *Arabidopsis*: *LEAFY* (*LFY*), *CAULIFLOWER* (*CAL*), *APETALA1* (*AP1*) and *FRUITFULL* (*FUL*) (Maizel and Weigel, 2004; Moyroud et al., 2001; Mandel et al., 1992).

After flower meristem specification, floral organ cell-fate determination occurs. The so-called ABC genes are necessary for this process (Figure 1a). Indeed, according to the ABC model of flower development the A genes (*APETALLA1 (AP1)* and *APETALA2 (AP2)*) are expressed alone in the outer whorl of the floral meristem and are necessary for sepal specification. A and B genes (*PISTILLATA (PI)* and *APETALA3 (AP3)*) are necessary for petal specification in the second whorl of the floral meristem, while B and C genes (*AGAMOUS (AG)*) together are necessary for stamen specification in the third whorl, and finally C alone is necessary for carpel specification (Coen and Meyerowitz, 1999) in the innermost whorl of the floral meristem (Stewart et al., 2016) (see Figure 1a). All of these genes, except *AP2*, are Type II MADS-box genes (Álvarez-Buylla et al., 2000) that codify for transcription factors with a DNA-binding domain (MADS), an intermediary domain (I), a putative protein-protein interaction domain (K) and a COOH putative transactivation domain (Coen and Meyerowitz, 1999; Ng and Yanofsky, 2001).

**Figure 1.**
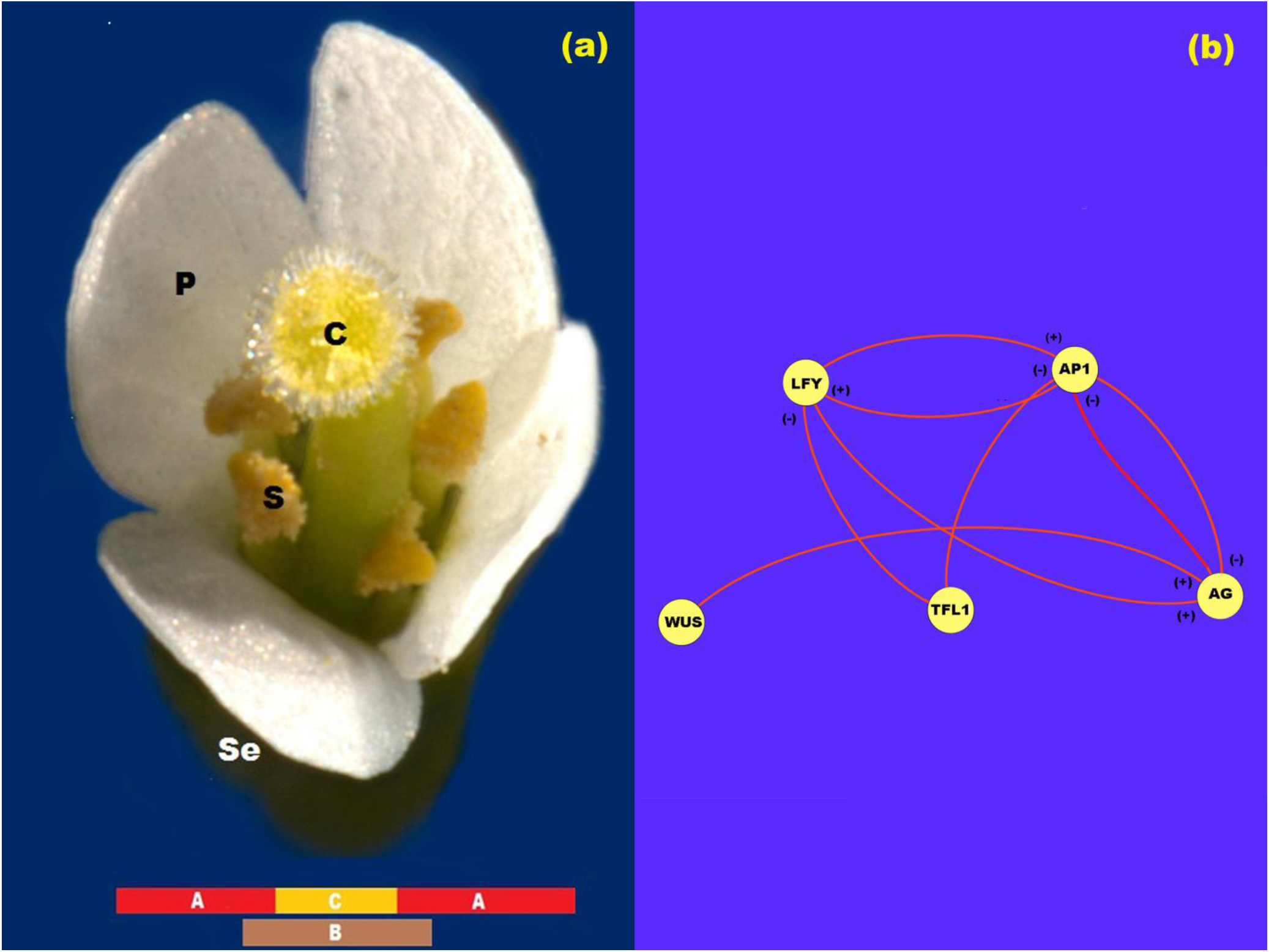
ABC model of flowering. a) ABC model of flowering for *Arabidopsis*. In this figure *se*: sepals; *p*: petals; *s*: stamen and *c*: carpel. b) Network representation of the interaction between the proteins LFY, AP1, TFL1, AG and WUS. In this Figure (+) represents activation and (-) represents inhibition.

The floral identity MADS-box genes *AP1* and *AG* have a central role in the ABC model. *AP1* is a direct target of the flowering time gene *FLOWERING LOCUS T* (*FT*) that responds to light inductive conditions and of *LFY* (Álvarez-Buylla et al., 2010). Upon formation of the flower primordia *AP1* is activated by LFY and by *FT* under long-day light inductive conditions and is expressed throughout the whole floral meristem (Pidkowich et al., 1999). Previous experiments have suggested that neither *AP1* mRNA nor AP1 protein move across the flower meristem (Sessions and Yanofski, 2000). *AG*, the C MADS-box gene, is activated by *WUS* (Espinosa-Soto et al., 2004; Jönsson et al., 2005; Jack, 2004; Ikeda et al., 2009). It has also been suggested that *WUS* is necessary to release the inhibitory effect of *AP1* over *AG*. Once *AG* is expressed, its protein represses *AP1* in the two central whorls, thus allowing for the spatial patterning of the floral meristem and the expression of the class B MADS-box genes (Jack, 2004).

Once the four whorls have been patterned, the AP1 protein forms complexes with a still unknown MADS-domain protein at the time of sepal identity specification in the first whorl, and AP1 interacts with APETALA3 (AP3), SEPALLATA (SEP) and PISTILLATA (PI) and this complex is necessary for petal specification in the second whorl. AG, in turn, interacts with SEP, PI and AP3 to form a protein quartet transcription complex required for stamen specification in the third whorl and finally AG associates with SEP genes to form the quartet transcriptional complex that is necessary for carpel specification in the fourth whorl (Pidkowich et al., 1999; Jack, 2004; Goto and Meyerowitz, 1994; Pelaz et al., 2000; Pelaz et al., 2001). Of relevance is the fact that *TERMINAL FLOWER1* (*TFL1*) counterbalances the action of floral meristem identity genes, *LFY, AP1* and *AG* (Parcy et al., 2002). *TFL1* encodes a protein that is highly similar to the animal RAF kinase inhibitors (Scheres, 1998). TFL1 specifies inflorescence meristem identity and induces the indeterminate nature of the inflorescence.

As data accumulate on the complex regulatory networks that underlie plant and animal development, it is becoming possible and necessary to postulate formal dynamic models. These may be now grounded on such data, and at the same time are useful to integrate necessary and sufficient regulatory modules for pattern formation and help uncover experimental holes. Such models hence constitute formal frameworks to test novel hypotheses *in silico* that can then be tested *in vivo*, and they are also the basis for understanding how spatio-temporal patterns of gene expression are established during development. Several regulatory network models for cell fate determination have been proposed (Espinosa-Soto et al., 2004; Álvarez-Buylla et al., 2008). These models describe the dynamics of the genetic network that sustain cell differentiation during flower development and they are mostly single-cell models.

The model proposed in Espinosa-Soto et al. (2004) uncovered what seems to be the core of a regulatory module that robustly converges to documented combinatorial gene activities characteristic of each floral organ primordia. In Espinosa-Soto et al. (2004), it is shown that a 15-gene regulatory dynamic network model that incorporates the ABC genes, as well as eleven non-ABC genes (Barrio et al., 2010) constitutes a regulatory module that robustly converges to 10 steady gene expression configurations that correspond to combinations of gene expression that have been experimentally documented for inflorescence and floral organ primordial cells. Four of these steady states correspond to a configuration of gene activation that characterize inflorescence meristem cells, while the other six attractors correspond to primordial cells of sepals (1), petals (2), stamens (2) and carpels. Four of the fifteen genes included in the floral organ specification network seem to be directly responsible for the spatio-temporal patterning of the floral meristem. These genes are *LEAFY* (*LFY*), *APETALA1* (*AP1*), *AGAMOUS* (*AG*) and *TERMINAL FLOWER1* (*TFL1*) (Álvarez-Buylla et al, 2010; Pidkowich et al., 1999; Jack, 2004; Parcy et al., 2002), but their mechanism of action during flower patterning is not clear.

Although GRN single-cell models has been successful to uncover the set of interactions that are both necessary and sufficient to recover the combinations of gene expression levels that characterize different primordial cells during early flower development in *Arabidopsis*, these models do not address how the spatio-temporal pattern of cell-fate determination is attained during flower development or what could be the role of transcription factors whose role is non-autonomous at the cellular level (Haspolat et al., 2019; Wang et al., 2014). In this direction, relatively few attempts have been done to understand the mechanisms underlying the emergence of spatiotemporal patterns (Jönsson et al., 2005; Dupoy et al., 2008; Alexeev et al., 2005; Barrio et al., 2010).

Some of such recent studies are suggesting that the emergence of spatiotemporal morphogenetic patterns partially depend on the uncovered intracellular regulatory networks (Álvarez-Buylla et al., 2008), but should also consider additional mechanisms that underlie the emergence of positional information. For example, in Barrio et al. (2010), a reduced version of the floral organ determination network was coupled with a physical field to explore the emergence of floral organ spatiotemporal patterns in wild type and mutant plants. In this work, the coupling of both fields leads to an interplay in which the macroscopic physical field breaks the symmetry of the floral meristem at any time, and gives rise to the differentiation of the meristem cells via a signal transduction mechanism that acts directly on the Gene Regulatory Network (GRN) that regulates cell-fate decisions during flowering.

In this direction, the works of Jönsson et al. (2005) and Gruel et al. (2016), propose a dynamic continuous system based on experimental results to study the underlying mechanism of *WUSCHEL (WUS)* spatial patterning during early stages of floral meristem determination and flower development (Alexeev et al., 2005). *WUS* is required for flowering and shoot and flower maintenance, it is stopped by *WUS* recessive mutations. In Alexeev et al. (2005), the authors proposed a reaction-diffusion model in which *WUS* is expressed in every point of the floral meristem unless a spatially distributed repressor signal is present. This repressor signal is induced by a signal from the extremes of the L1 sheet, and restricts *WUS* expression to the center of the sheet. The model accurately reproduces experimental observations in a two dimensional lattice of cells, and relates the repressor signals to CLAVATA3 (CVL3) signaling. However, recovered patterns are not robust to variations in the parameters. Similar results were obtained by Gruel et al. (2016) who showed that the combination of signals originating from the epidermal cell layer, which include the CVL3-WUS negative feedback loop, can correctly pattern gene expression domains.

Thereby, the present contribution further elaborates on previous spatio-temporal models and explores the emergence of the four whorls of differential gene expression in the L1 layer of floral meristem cells in concordance with the ABC model of flower patterning. Our model shows how the four-whorl symmetry of the floral meristem dynamically arises from a spatially homogenous distribution of expression of *LFY, TFL1, AP1, AG* and *WUS* (Espinosa-Soto et al., 2004). The model takes into account the nonlinear interactions between AP1, AG, LFY and TFL1 proteins during early flower development, and it also includes the equations for the spatial patterning of *WUS* expression presented in the work of Alexeev et al., (2005). We postulate that *WUSCHEL* spatial pre-pattern of expression *is a necessary but not sufficient* condition for the patterning of the floral meristem into the four whorls. *WUS* pre-pattern breaks the initial symmetry of the system and induces the expression of *AG* in the third and fourth whorls, and gives rise to a new symmetry that corresponds to the ABC model of gene expression Gruel at al. (2016).

The model also tests the role of *LFY* during the patterning of the floral meristem. *LFY* is a meristem-identity gene that responds to several internal and external flowering-inducing signals and also has a central role in regulating the patterns of the ABC genes (Álvarez-Buylla et al., 2008). At the same time, this gene is regulated for example by the flowering time gene *SUPPRESSOR OF OVEREXPRESSION OF CONSTANS* (*SOC1*) gene that integrates the flowering response to light, vernalization and gibberellins (GA), and is also a direct target of GA (Álvarez-Buylla et al., 2010; Pidkowich et al., 1999; Scheres, 1998; Villarreal et al., 2012; Boss et al., 2004; Okamuro et al., 1996; Traas and Vernoux, 2002). Previous experimental work has provided evidence for the movement of LFY protein, from the L1 layer into the internal layers L2 and L3 of the apical meristem, during flower development (Ingram, 2004). Thus, LFY forms a gradient of activation that extends from the L1 to the L3 sheet of the SAM (Wu et al., 2003). Experiments carried out with the reporter Green Fluorescent Protein (GFP) expressed under the action of the *LFY* promoter have shown that the protein LFY moves along the L1 sheet of the SAM, where it forms a uniform field of activation (Wu et al., 2003). These results suggest that *diffusion of this protein is probably not critical for the spatial patterning of the L1 sheet during floral organ primordia specification* but no dynamic mechanism had been proposed for this. In the context of the model presented here, we show that the movement of LFY *along* the L1 sheet of the floral meristem is not a necessary condition for the onset of the ABC pattern of gene expression.

In conclusion, the aim of the model presented in this work is to demonstrate that the interaction of the four chemical fields generated by the interaction of LFY, TFL1, AG, AP1 and WUS can pattern the L1 cell layer into the three domains of gene expression according to the ABC model of flowering. The model suggests five main points: a) LFY diffusion does not take a fundamental part in the patterning of the floral meristem *along* the L1 sheet of cells; b) the pattern obtained from the model defines three domains of gene expression according to the ABC model of flowering; c) WUS pre-pattern *is a necessary but not a sufficient condition* for the correct patterning of the L1 layer of the floral meristem; d) the spatio-temporal distribution of *LFY, AP*1, *AG*, and *TFL*1 products along the L1 sheet can effectively be a necessary but not sufficient condition for floral organ determination, once the WUS pre-pattern has been established; e) exists, at least, a set of parameters values for which we can obtain a solution of the model that resembles the experimentally observed ABC pattern.

## 2 Model

In the model, we propose hypothetical 15 cells along the L1 layer of the floral meristem with a near uniform average size of about 4.4 µm each one. In consequence, the diameter of the layer is ∼ 66 µm. We assume that each one of these ∼ 15 cells along the diameter of the meristem is characterized only by the amount of the protein produced by *LFY, AP1, AG, WUS*, and *TFL1* at time *t*, which is a measure of the activation level of the respective gene. In the model, we covered the L1 layer with 15 of these idealized cells.

In order to test only the role of the interaction of these proteins in the patterning of the L1 sheet, we assume that during the time of simulation the size of the L1 layer is constant and that the LFY difference of concentration along the L1-L3 direction is small enough to no significantly affect LFY concentration in the L1 sheet during the time of simulation.

In the research papers of Espinoza-Soto et al. (2004), Álvarez-Buylla et al. (2008), Barrio et al. (2010), and Villarreal et al. (2012), the experimental gene data that support the regulatory interactions of *LFY, AP1, AG*, and *TFL1* during floral induction are summarized and formalized in the form of tables of logical rules. The mathematical model presented below is a direct translation of these logical rules into its corresponding continuous mathematical expressions (Figure 1b). Thus, the logical rules are used as a guidance to establish the equations that are postulated here to drive the ABC patterning process. In these mathematical equations we represent the amount of each protein with their respective name in lower case italic letters.

In this form, from Figure 1b we propose that the rate of *LFY* activation results from a balance between the intrinsic rate of activation of the gene (*k*_*1*_), the rate at which it is activated by protein AP1, the rate at which it is inactivated by protein TFL1 and the intrinsic rate of inactivation of the gene itself. Finally, we must take into account the interaction among L1 cells due to LFY movement. According to the method of discretization of the meristem we obtain the equation:

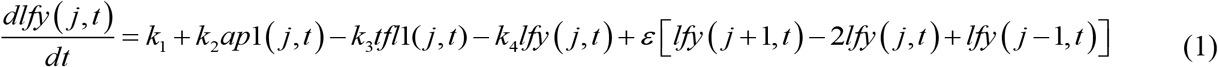

where *j* = 1, 2, 3,…, 15 is the number of the cell, 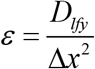 is the coupling coefficient between cells, *D*_*lfy*_ is the diffusion coefficient of LFY and Δ*x* is the length of a idealized cell. Protein LFY cannot flow out of the meristem though the extremes of the array of cells, and is initially distributed at a uniform basal concentration along it.

From Figure 1b, the rate of *AP1* activation results from a balance between its intrinsic rate of activation (*k*_*5*_), the rate at which it is activated by LFY protein, the rate at which it is inactivated by TFL1 protein, and the rate of inactivation of the gene itself. Once the *AG* gene is activated as a result of the presence of WUS protein in the centre of the flower meristem, AG protein turns off *AP1* activity from the zone corresponding to the third and fourth whorls and AP1 protein turns off *AG* activity from the first and second whorls. As we mentioned before, neither AP1 nor AG seem to diffuse among cells. Thus, the spatial patterning of the L1 cell layer of the presumptive floral meristem lies on the exclusion action between these two proteins by a yet unknown kinetic mechanism. Consequently we propose the following equations that describe the activation of *AP1* in cell *j* at time *t*:

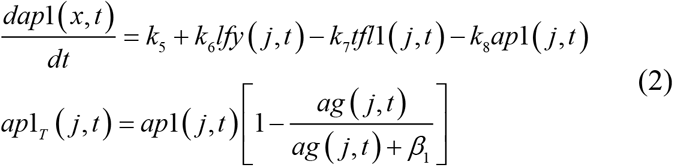

where *ap1*_*T*_(*j,t*) is the distribution of AP1 protein along the meristem due to the presence of AG protein.

As reviewed in Espinoza-Soto et al. (2004) and Goto and Meyerowitz (1994), the rate at which *AG* is activated depends on its rate of activation by LFY protein, the rate at which it is inactivated by TFL1 protein and its rate of inactivation. The rate at which *AG* activation level increases in the system tightly depends on the WUS protein pre-pattern (Figure 1b). According to Álvarez-Buylla et al., (2010) and Espinosa-Soto et al. (2004) there is a double negative loop between *AP1* and *AG*, in which *AG* inhibits *AP1* expression from whorls 3 and 4, and *AP1* inhibits *AG* expression from whorl 1 and 2. In this form, we propose a noncompetitive inhibition of AP1 protein on the production of AG:

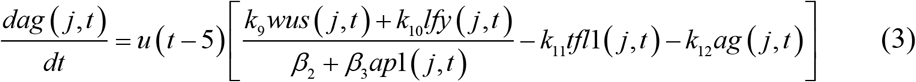

where *u*(*t* – 5) represents the unitary step function that lags AG spatial pattern formation until *t* = 5 h. We are not explicitly modeling the mechanism that regulates flowering time and the function *u* is necessary for the correct timing of the process in the model. However, if *u* is not used the AG spatial pattern emerges after a few integration steps. In every case, *AG* spatial expression pattern arises once the *WUS* expression pre-pattern is established.

As reviewed in Álvarez-Buylla et al., (2010) and Espinosa-Soto et al. (2004), the rate at which *TFL1* activation level increases in the system results from a balance between its intrinsic rate of activation (*k*_*13*_), the rate at which it is inactivated by LFY protein, the rate at which it is inactivated by AP1 protein and its rate of inactivation:

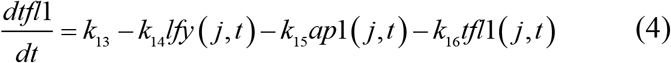

Jönsson et al. (2005) shown that the pattern of *WUS* expression has its maximum approximately at the center of the L1 fourth whorl, and does not expand too far from this center (Figure 2a). In this work, we adapted the repressor model of Jönsson et al. (2005), which consists of the following equations:

**Figure 2.**
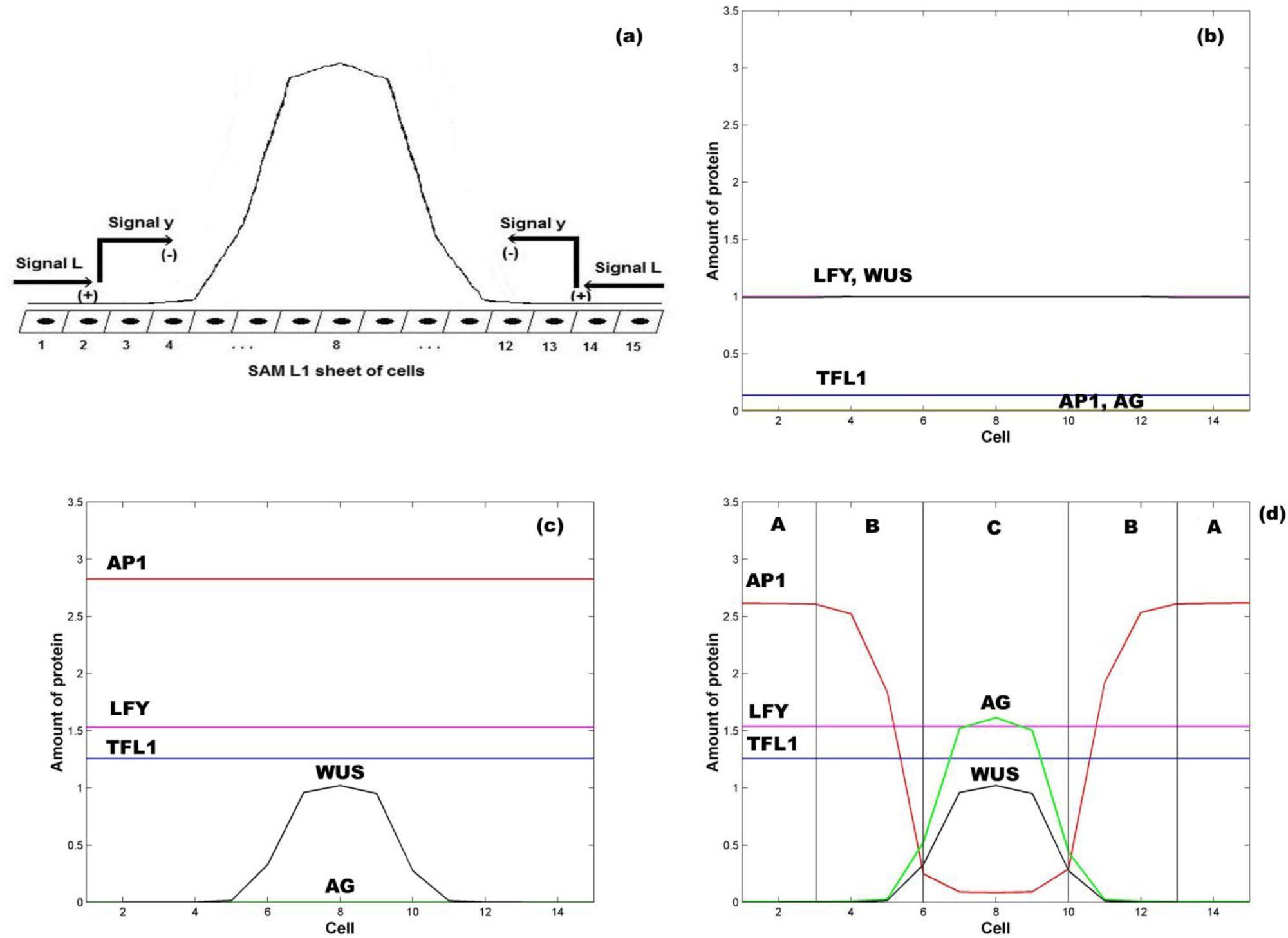
Emergence of the ABC zones of flower organ determination. a) WUS pre-pattern is the result of the action of the inhibitory signal *L* from the extremes of the SAM L1 sheet that induces the activation of the inhibitory chemical signal *y* that restricts *WUS* expression to the inner whorl of the floral meristem. In the model we represent the floral meristem as a linear array of 15 cells that crosses the diameter of the four whorls. b) Initial homogeneous spatial distribution of the chemical fields at the beginning of the simulation, LFY (red line), TFL1 (yellow line), AP1 (brown line) AG (black line) and WUS (blue line); c) WUS pattern (blue line) arises at the center of the floral meristem after ∼ 1 h; d) the initial homogenous state of the floral meristem is completely broken after ∼ 16 hours. AG is expressed at the center of the meristem (black line) and its presence moves AP1 away from this zone. In consequence, the floral meristem has been patterned into three well defined zones of gene expression. In all Figures *ε* = 5. In all panels *L*(1) = *L*(2) = *L*(3) = *L*(13) = *L*(14) = *L* (15) = 1, and *L*(*j*) = 0 for 4 ≤ *j* ≤ 12; in similar form: *y*(1) = *y*(2) = *y*(3) = *y*(13) = *y*(14) = *y* (15) = 1 and *y*(*j*) = 0 for 4 ≤ *j* ≤ 12.

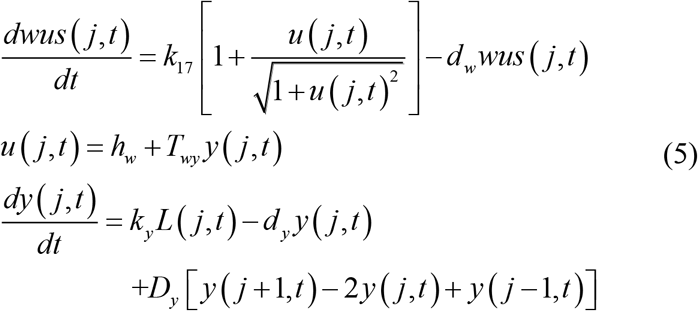

subject to the following boundary conditions:

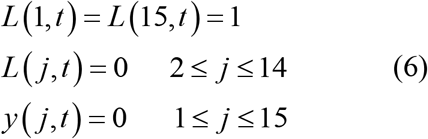

The model was solved using the Euler predictor-corrector method. The simulation was done for 1,200,000 time steps of 0.05s which represents 16.6 hrs. The initial condition used in this work are: *lfy*(*j*,0) = 1, *ap*1(*j*,0) = 0, *ag*(*j*,0) = 0, *tfl*1(*j*,0) = 0.1 and *wus*(*j*,0) = 1 for *j = 1, 2, 3, …, 15*. Additionally: *y*(1,0) = *y*(2,0) = *y*(3,0) = *y*(13,0) = *y*(14, 0) = *y*(15,0) = 1 and *y*(*j*,0) =0 for *j = 4, 5, 6, …, 12; L*(1,0) = *L*(2,0) = *L*(3,0) = *L*(13,0) = *L*(14, 0) = *L*(15,0) = 1 and *L*(*j*,0) =0 for *j = 4, 5, 6, …, 12*.

In Table 1 we show the parameter values used in the model. We made parameter estimation by randomly varying each individual parameter value reported in the second column of Table 1 in a range of about ± 10% of its original value, and choosing those interval of values for which the model output is stable. These intervals of values are presented in the third column of Table 1.

**Table 1.** Parameter values for the spatial ABC patterning model of flowering.

| Parameter | Value in the Model | Interval of Parameter Values |
| --- | --- | --- |
| $k_1$ | $0.03 \mu\text{M s}^{-1}$ | [0.03, 0.035] |
| $k_2$ | $0.02 \text{ s}^{-1}$ | [0.02, 0.023] |
| $k_3$ | $0.02 \text{ s}^{-1}$ | [0.015, 0.02] |
| $k_4$ | $0.04 \text{ s}^{-1}$ | [0.035, 0.04] |
| $k_5$ | $0.09 \mu\text{M s}^{-1}$ | [0.9, 1.5] |
| $k_6$ | $0.05 \text{ s}^{-1}$ | [0.05, 0.07] |
| $k_7$ | $0.02 \text{ s}^{-1}$ | [0.01, 0.02] |
| $k_8$ | $0.05 \text{ s}^{-1}$ | [0.04, 0.05] |
| $k_9$ | $0.08 \text{ s}^{-1}$ | [0.08, 0.5] |
| $k_{10}$ | $0.025 \text{ s}^{-1}$ | [0.025, 0.05] |
| $k_{11}$ | $0.03 \text{ s}^{-1}$ | [0.01, 0.03] |
| $k_{12}$ | $0.05 \text{ s}^{-1}$ | [0.01, 0.05] |
| $k_{13}$ | $0.9 \mu\text{M s}^{-1}$ | [0.7, 0.9] |
| $k_{14}$ | $0.08 \text{ s}^{-1}$ | [0.07, 0.08] |
| $k_{15}$ | $0.03 \text{ s}^{-1}$ | [0.03, 0.08] |
| $k_{16}$ | $0.55 \text{ s}^{-1}$ | [0.55, 0.75] |
| $k_{17}$ | $0.05 \mu\text{M s}^{-1}$ | constant value |
| $\beta_1$ | $0.05 \mu\text{M}$ | constant value |
| $\beta_2$ | $1 \mu\text{M}$ | constant value |
| $\beta_3$ | 0.55 | constant value |
| $d_w, h_w, T_{wy}, k_y, d_y, D_y$ | 1.75, 2, -30, 0.2, 2, 0.1 | Jönsson et al. (2005) |

## 3 Results

The numerical integration of the set of equations postulated in the model leads to the results shown in Figure 2. In Figures 2a and 2b it is clear that the first genes that are switched *on* are *LFY* and *TFL1*. The activation level of these two genes is uniform along the presumptive floral meristem. As expected, LFY >>TFL1 at all times (see Table of Logical Rules in Espinosa-Soto et al., 2004) as required for floral induction.

Flower induction depends on numerous genes (∼ 2000) that respond to light, and to external and internal signals. However, *LFY* and *AP1* are two of the most important downstream targets of flower meristem specification and are key markers of flower meristem identity (Pidkowich et al., 1999; Jack, 2004; Boss et al., 2004). As we show in Figure 2c, before the new spatial pattern of the system is established, *AP1* is uniformly activated along the L1 cell layer, in response to *LFY* activation (Equation 2). *WUS* is activated in the center of the L1 cell layer under the action of an inhibitory signal *L* from the extremes of the layer (Jönsson et al., 2005).

In the model, *AP1* should be activated before *AG*, and the WUS pre-pattern must induce *AG* activation prior to *AP1* inhibition by AG in order to obtain the complete set of flower structures. In this form we obtain the sequence of events of gene activation): *LFY, AP1, AG* (Figure 2b, Figure 2c and Figure 2d) (Pidkowich et al,1999). *TFL1* is turned *on* at the same time that *LFY* comes *on* and remains at a low and homogeneous level of activation throughout early stages of flower development (Figure 2d) (Espinosa-Soto et al., 2004).

*WUS* expression in the flower center blocks the inhibitory effect of *AP1* over *AG*, allowing the expression of the latter in this field centered at ∼ cell 8 (Espinosa-Soto et al., 2004). *AG* is expressed in this field and exerts an increasing inhibitory effect on *AP1* as *AG* relative level of expression increases, according to Equation 2. Thus, these results from the model show that *this interplay, at the cellular level, given the WUS spatial pattern of activation in the flower center, is a necessary but not sufficient condition for the spatial patterning of the L1 cell layer of the SAM during the floral induction process*. As a result, this mechanism produces the expression of the class C MADS genes in the fourth whorl and the class A MADS-box genes in the first whorl. Class B genes are expressed in the cells between these two peaks of opposite activity (Figure 2d).

*WUS* pattern is due to the inhibitory signal *L* from the cells of the extreme of the L1 layer. Figure 2d is obtained when the signal *L* is present in cells 1, 2, 3, 13, 14 and 15. When the signal *L* is reduced to cell 1 in the left extreme, and to cell 15 in the right extreme (*L*(*j,t*) = 1 for *j* = 1, 15 and *L*(*j,t*) = 0 for 1 < *j* < 15) the qualitative form of the pattern shown in Figure 2d is conserved, but it becomes broader and asymmetric with respect to cell 8 (Figure 3). This numerical result indicates that the signal *L* is the primary factor that patterns the extent of the spatial expression of the *WUS* and *AG* genes, and breaks the initial system symmetry through the set up of a diffusible inhibitory signal *y* that is initially presented only in the extremes of the L1 cell layer (Jönsson et al., 2005) (Figure 2a). The molecular identity of the *L* and *y* signals still remains unclear (Jönsson et al., 2005). However, one possibility is that these inhibitory signals could be diffusible peptides of the CLV family (Alexeev et al., 2005; Sablowsky, 2009; Gruel et al., 2016). It is possible that the fields of mechanic and elastic forces also underlie positional information important for spatial patterning (see Barrio et al., 2010).

**Figure 3.**
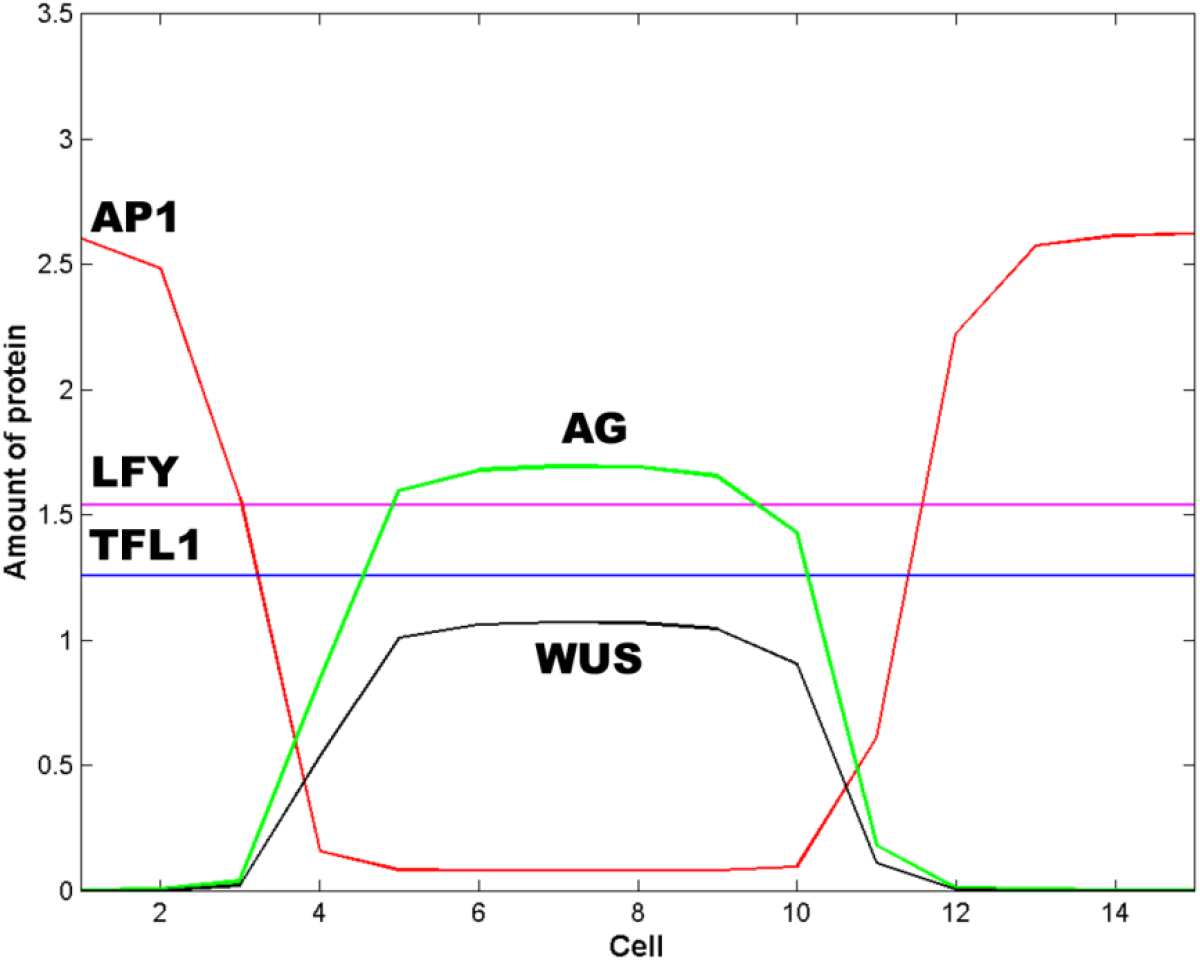
Effect of the spatial extent of the inhibitory signals *L and y*. In this Figure *L* = 1 and *y* = 1 for cells 1 and 15; *L* = 0 and *y* = 0 otherwise. The effect of decrease the spatial extent of the inhibitory signals *L* and *y* is to pattern the floral meristem into a spatio-temporal stable dissipative structure, which becomes broader and asymmetric with respect to cell 8 and resembles an altered floral structure. In this Figure *t* = 16 h and *ε* = 5.

In Figure 2d we show the state of each of the 15 cells of the model at steady state conditions after the spatial patterning process of the presumptive floral meristem. As shown in Figure 1, the formation of floral structures depends on the correct set up of the four zones of gene expression configurations (Álvarez-Buylla et al., 2010). Our model renders a spatio-temporal patterns of gene expression with a clearly defined A zone at the outer whorl, and a C zone of expression centered at the fourth whorl. The B zone lies between these two zones overlapping with A in the second and with C in the third whorls (Figures 2d and 3). This pattern mimics that found during early stages of *Arabidopsis* flower development, and we should remark that the entire dynamics of the system rests on the boundary conditions set at the extremes of the modeled domain of cells (see above paragraph).

Zone A is characterized by high levels of expression of *LFY* and *AP1*, and a low level of *TFL1* expression. Zone C has high levels of *WUS, AG* and *LFY* expression and low *TFL1* expression levels. Zone B has a combination of different levels of expression of the five genes. In this form, in each zone the complete network of 15 genes coupled to the continuous signal fields modeled here yields a spatio-temporal pattern that mimics that observed during early flower development (Espinosa-Soto et al., 2004). The minimal network modeled here is also useful to address the role of the intercellular movement of LFY that is a key factor during flower development (Figures 2d and 3).

Protein LFY can move among cells along the L1 cell layer (Wu et al, 2003). If we vary the coupling factor *ε* from 0 to a value of 10, we do not observe any change in the recovered spatial or temporal patterns concerning the level of expression of *LFY* itself, and also of *TLF1, AP1* and *AG*. This result suggests that free diffusion of LFY *among* cells is not critical for the observed spatial patterning of the key regulatory genes involved in early flower development (Wu et al, 2003), but LFY is the chemical force that drives the reaction processes that induce the instability of the chemical field during the symmetry breaking process (Equations 1-3, Figure 1b).

In order to further address the role of LFY diffusion in sustaining the steady state dissipative structure formed after the spatial patterning of the system emerges, we made a series of simulations in which *ε* was varied randomly every 50 s, the final dissipative structure is not altered, indicating that the interactions responsible for the preservation of this structure are independent of the flux of LFY between cells down the L1 layer. Furthermore, if we allow random values *of ε* among L1 cells the system evolves to the same dissipative structure. These results support the idea that *the role of LFY in the spatial patterning process of L1 during flower development does not depend on its diffusive properties but on its flower meristem identity function in interaction with several other components of the flower organ specification GRN, including its regulatory interactions with the ABC genes, and in response to several inductive factors* (Pidkowich et al., 1999; Jack, 2004; Scheres, 1998).

## 4 Discussion

Reaction-diffusion processes have been shown to be important components of the mechanisms underlying the emergence of ordered spatio-temporal patterns of gene expression patterns in biological systems. The pioneer work of Turing (1952), and the posterior works of Prigogini and Nicolis (1967), Prigogine and Lefever (1968), and Gierer and Meinhardt (1972), have shown that chemical dissipative structures form fields that are a source of positional information (Wolpert, 1994). However, it is no clear yet how this positional information is interpreted by gene networks; although some attempts have been done in this direction in the case of animal systems (Currie and Ingham, 1998; Jaeger et al., 2004).

In the particular case of *Arabidopsis* flower development, recent works have tried to link the Boolean dynamics of the genetic network for floral determination proposed by Espinosa-Soto et al. (2004), with the ABC model of flower development. However, the ABC model does not provide a dynamical explanation for the emergence and maintenance of the steady-state spatial patterns of gene expression that characterize each primordial floral organ cell type as a result of ABC and non-ABC gene interactions.

Espinosa-Soto et al. (2004), proposed a discrete dynamic model of the necessary and sufficient set of ABC and non-ABC genes interactions to recover the gene configurations that are characteristic of the four floral organ cell-fates. This model postulates a network of interaction among 15 genes (nodes). The model shows that all possible initial conditions lead the system to a few steady states of gene activity that match the gene expression profiles observed in four regions of the inflorescence meristem (with neither *UFO* or *WUS*, with both or either one of these two factors), and in each of the four types of floral organ primordial cells. A conclusion from this model is that floral cell fate determination is determined by the structure and dynamics of the GRN proposed, which can be considered as a robust developmental module underlying cell-fate determination during early stages of flower development. This model cannot be used to address the mechanisms underlying the emergence of positional information and the spatiotemporal patterns during flower development.

A stochastic version of the dynamics of the gene network proposed by Espinosa-Soto et al. (2004), to explore cell-type transitions is presented by Álvarez-Buylla et al. (2008). Although the basic dynamical features of the network remain Boolean, the introduction of different uncertainty levels in the updating of the logical rules mimics the effect of noise on the GRN that can be due to external fluctuations or internal noise due to sampling errors in the transcription factors involved. The model exhibits recovers the temporal pattern of cell-fate transitions observed during flower development, but does not include a spatially explicit domain.

In order to explore the emergence of positional information and spatial patterning during flower development, the Boolean dynamics of the GRN proposed by Espinosa-Soto et al. (2004), is coupled to elastic fields in the floral primordium (Barrio et al., 2010). The main hypothesis in this work is that there is at least one mechanical field that breaks the symmetry of the floral primordium at a given time during early stages of flower development. This field provides the positional information required for the process of cell differentiation in different spatial domains of the primordium as a result of the dynamical coupling via a signal transduction mechanism that, in turn, acts directly upon the gene regulatory network underlying cell-fate decisions within cells. It is then the feedback between the intracellular GRN and such extra-cellular signals and fields that underlies positional information and spatial patterning. This model is able to recover the multi-gene configurations characteristic of sepal, petal, stamen, and carpel primordial cells arranged in concentric rings, in a similar pattern to that observed during actual floral organ determination. An important caveat of this model is that it assumes the existence of a field *ϕ* that *a priori* breaks the symmetry of the floral meristem. The model is a hybrid one, in which the equations of the mechanical field are continuous, and the states of the GRN are discrete.

A general theory for genotype to phenotype mapping is proposed by Villarreal et al. (2012). In this work the authors have put forward an analytical derivation of the probabilistic epigenetic landscape for an N-dimensional genetic regulatory network grounded on experimental data. This method was applied to the *Arabidopsis thaliana* floral organ specification GRN used in Espinosa-Soto et al., (2004) successfully recovering the steady-state gene configurations characteristic of primordial cells of each floral organ type in wild-type and ABC mutants, as well as their temporal patterns of transitions that mimics that observed in actual flower development when ABC gene decay rates are relatively similar to those which have been reported experimentally.

Some of the previous modeling approaches have attempted to integrate the GRN underlying floral organ specification with coupling mechanisms that recover observed spatial patterns during early flower development. An additional effort to model the mechanisms underlying floral organ specification is presented in Wang et al. (2014). In this paper, authors use a continuous approach and specifically consider the dynamical response of *AP1* and *LFY* to photoperiod.

Previous studies have shown, using flower development as study system, that *the structure and dynamics of the floral organ specification GRN underlies the attractors attained during its temporal evolution*, and that the kinetic rates of interaction between their nodes are important for determining the timing and responsiveness of the GRN being considered. Furthermore, additional studies have shown that the spatial interactions among cells through short or large-range diffusible signals is a necessary condition for the emergence of dissipative structures in any multi-cellular system with nonlinear dynamics (Prigogine and Nicolis, 1967). In this study we have explored the link between the GRN dynamics and the emergence of apical meristem regions with specific positional information that had remained unclear from previous studies.

We explored how the nonlinear interaction between the protein products of the floral gene regulatory network yields the instability of the chemical fields in the flower primordium, and how the diffusive properties of some of these proteins drive the system into a steady stable dissipative structure with a pattern that coincides with that observed during floral organ specification in early flower development.

Hence, we proposed without *a priori* assumptions concerning the symmetry of the L1 sheet of cells, *that the subnet of five nodes WUS, AP1, AG, LFY, and TFL1, comprise a minimal GRN necessary for the initial patterning of the floral meristem* (Figures 2d and 3). The necessary condition for the patterning of the floral meristem into the A, B and C zones is the pre-patterns of WUS. The dynamical properties of this net are determined by the kinetic parameters of the strength and timing of the interactions among nodes, and by the diffusive properties of LFY and the inhibitory signal *y*.

In our work, the molecular interactions that determine floral organ induction are modeled with a set of coupled nonlinear differential equations, while the interaction among the L1 sheet of cells, due to the diffusion of LFY and signal *y*, is modeled with the discrete version of the Laplacian. The intensity of the coupling among the floral meristem cells is determined by the values of the coupling coefficients *ε* and *Dy* (See Model section).

Our model seeks to elucidate how the nonlinear interaction between the protein products of WUS, *LFY, TFL1, AG* and *AP1* may be involved in patterning the floral meristem and if such minimal GRN is sufficient to achieve so. For this purpose we used a linear arrange of 15 cells that extends along the diameter of the four whorls and we initialize our simulations by setting homogeneous initial conditions for all the cells of this array (Figure 2b). We couple this homogeneous chemical field to the reaction-diffusion process that produces the WUS spatial pre-pattern centered at whorl 4 (Jönsson et al., 2005) (Equation 5). In the work of Jönsson et al. (2005) the forces that pattern WUS spatial distribution are taken as unknown signals *L* and *y* from the extremes of the L1 sheet. In the work of Alexeev et al. (2005) it is suggested that at least one of the unknown signals could correspond to the negative regulatory effect that CLV3 has over WUS spatial distribution. The second inhibitory signal could be AG, which has been demonstrated to negatively regulate *WUS* spatial pattern of expression (Liu et al., 2011).

As we mentioned before, LFY has diffusive properties that could take part in the definition of the ABC zones. However, as we show in the Results section, random variations in the coupling coefficient *ε* (see Results section) that stands for intercellular LFY movement along the L1 sheet does not affect the final spatial pattern of the system. This result suggests that LFY diffusion is not necessary for the spatial patterning of A, B and C functions in the L1 layer. In this form, the entire spatial dynamics depends on the diffusion of the inhibitory signals *L* and y discussed above (see Figure 3). Moreover, the numerical solution of the model shows that, for the particular set of parameters values shown in Table 1, WUS pre-pattern is a *necessary but not sufficient condition* for the patterning of the floral meristem into the four spatially distributed chemical fields postulated by the ABC model.

The model reproduces the initial sequence of events during floral organ specification. This sequence is formed by an initial expression of the genes *AP1, LFY* and *TFL1* in all cells (Figure 2b), followed by the emergence of the *WUS* pattern. The regional activation of *WUS* centered at the fourth whorl breaks the homogeneity of the initial chemical field of the system (Figures 2b and 2c). Once the WUS pattern is formed, *AG* is expressed and exerts its inhibitory action on *AP1* in the center of the cell array, fixing *AP1* expression at the extremes (first whorl) of the floral meristem (Figure 2d). In order to obtain the correct qualitative pattern of floral induction, it is necessary to take into account the mutual inhibition loop formed by *AP1* and *AG* (Espinosa-Soto et al., 2004). Furthermore, this loop seems to be necessary for the stability of the pattern (see Results section).

Experimental data indicates that *WUS* excludes *AP1* expression from the fourth whorl and thus activates *AG*. The model assumes that *AG* is activated prior to AP1 exclusion from the fourth whorl. But if the AP1 exclusion function (Equation 2) of the model is written in terms of WUS instead of AG, the qualitative form of the final pattern of floral organ induction is not altered, indicating that the patterning of the system does not depend if either *WUS* and *AG* genes exerts the inhibitory action over *AP1*. However, the floral organ specification GRN proposed in Espinosa-Soto et al. (2004), states that is *AG* who inhibits *AP1*.

In this form, from the numerical solution of our model it is possible to obtain a chemical dissipative structure that patterns the linear array of 15 L1 cells into three well defined zones of differential expression of the five genes of the subnet modeled here. Each zone (whorl) has positional information that is interpreted in the form of a specific combination of the A, B and C genes that coincides with the necessary conditions for organ determination in each whorl as postulated by the ABC model.

Finally, it is important to mention that in this work we did not perform ABC mutant simulations because we used a subnet of only five of the 15 nodes of the floral organ specification GRN proposed before (Espinosa-Soto et al., 2004; Barrio et al., 2010). The interaction of these five nodes with the rest is important to recover the floral patterns observed in mutant plants.

## 5 Conclusions

The aim of our computational model is to propose a probable mechanism for the spatial patterning process of the presumptive floral meristem based on the mutual exclusive interaction at a cellular level of the AP1 and AG, and a spatial pre-pattern of WUS (Jönsson et al., 2005) centered at the fourth whorl, which is a necessary but not sufficient condition for floral organ determination. Our model has also enabled us to show that although experiments with *LFY*:GFP hybrids clearly show that LFY can effectively move from cell to cell along the L1 sheet of cells of the SAM (Wu et al., 2003), LFY diffusion has no effect on the onset or maintenance of the peaks of *AP1* and *AG* activity predicted by the model, which mimic the ABC patterns.

The dissipative structure obtained from the numerical solution of the model shows two opposite peaks of activity at the first and fourth whorls formed by AP1 and AG, respectively, that define the A and C zones of floral induction. The B zone lies in the middle of these peaks and represents different combination of expression of the five genes in whorls 2 and 3. Thus, the numerical solution of the model proposed in this work leads to the onset of the four chemical fields that contain the positional information required for the differential expression of the A, B, and C genes according to the ABC model for floral organ specification. These four coupled chemical fields form a dissipative structure that resembles the floral organization observed during the early stages of development in the floral primordium.

Finally, the model presented in this work suggest five main points susceptible to be experimentally tested: a) LFY diffusion does not take a fundamental part in the patterning of the floral meristem *along* the L1 sheet of cells; b) the pattern obtained from the model defines the ABC zones of gene expression according to the ABC model of flowering; c) WUS pre-pattern *is a necessary but not a sufficient condition* for the correct patterning of the L1 layer of the floral meristem; d) the spatio-temporal distribution of *LFY, AP*1, *AG*, and *TFL*1 products along the L1 sheet can effectively be a necessary but not sufficient condition for floral organ determination, once the WUS pre-pattern has been established; e) exists, at least, a set of parameters values for which we can obtain a solution of the model that resembles the experimentally observed ABC pattern.

## 7 Data Availability Statement

The original contributions presented in the study are included in the article/supplementary material; further inquiries can be directed to the corresponding author.

## 8 Author Contributions

Both authors made equal substantial contributions to this manuscript.

## 10 Acknowledgments

We thank Erika Juárez and Diana Romo for technical and logistical assistance. We also thank Yamel Ugartechea for Figure 1.

## References

Alexeev, D.V., Ezhova, T.A., Kozloy V.N., Kudryaytsev, V.B., Nosov, M.V., Penin, A.A., Skryabin, K.G., Choob, V.V., Shulga, O.A., and Shestakov, S.V. (2005). Spatial pattern formation in the flower of Arabidopsis thaliana: mathematical modeling. Doklady Biological Sciences 401:133–135.

Alvarez-Buylla, E.R., Chaos, A., Aldana, M., Benítez, M., Cortes-Poza, Y., Espinosa-Soto, C., Hartasánchez, D.A., Lotto, R.B., Malkin, D., Escalera-Santos, G.J., and Padilla-Longoria, P. (2008). Floral morphogenesis: stochastic explorations of a gene network epigenetic landscape. PLoS ONE 3: e3626. doi: 10.1371/journal.pone.0003626

Alvarez-Buylla, E.R., Liljegren, S.J., Pelaz, S., Gold, S.E., Burgeff, C., Dittal, G.S., Vergara-Silva, F., and Yanofsky, M.F. (2000). MADS-box gene evolution beyond flowers: expression in pollen, endosperm, guard cells, roots and trichomes. The Plant Journal 24:457–466. doi: 10.1046/j.1365-313x.2000.00891.x.

Alvarez-Buylla, E.R., Benítez, M., Corvera-Poiré, A., Chaos, A., de Foltier, S., Gamboa de Buen, A., Garay-Arroyo, A., García-Ponce, B., Jaimes-Miranda, F., Pérez-Ruiz, R.V., Piñeyro-Nelson, A., and Sánchez-Corralesa, Y.E. (2010). Flower development. The Arabidopsis Book 8:e0127

Barrio, R.A., Hernandez-Machado, A., Varea, C., Romero-Arias, J., Alvarez-Buylla, E.R. (2010). Flower development as an interplay between dynamical physical fields and genetic networks. PLoS ONE 5:e13523. doi:10.1371/journal.pone.0013523

Boss, P.K., Bastow, R.M., Mylne, J.S., and Dean, C. (2004). Multiple Pathways in the Decision to Flower: Enabling, Promoting, and Resetting. The Plant Cell 16:S18–S31. doi: https://doi.org/10.1105/tpc.015958

Coen, E.S., and Meyerowitz, E.M. (1999). The war of whorls: genetic interactions controlling flower development. Nature 353:31–37. doi: 10.1038/353031a0

Currie, P.D., and Ingham, P.W. (1998). The generation and interpretation of positional information within the vertebrate myotome. Mechanisms of Development 73:3–21. doi: 10.1016/s0925-4773(98)00036-7

Dupoy, L., Mackenzie, J., Rudge, T., and Haseloff, J. (2008). A System for Modelling Cell –Cell Interactions during Plant Morphogenesis. Annals of Botany 101:1255–1265. doi: https://doi.org/10.1093/aob/mcm235

Durfee, T., Roe, J.L., Sessions, R.A., Inouye, C., Serikawa, K., Feldmann, A., Weigel, D., and Zambryski, P.C. (2003). The F-box-containing protein UFO and AGAMOUS participate in antagonistic pathways governing early petal development in Arabidopsis. PNAS 100:8571–8576. doi: 10.1073/pnas.1033043100

Espinosa-Soto, C., Padilla-Longoria, P., and Alvarez-Buylla, E.R. (2004). A gene regulatory network model for cell-fate determination during Arabidopsis thaliana flower development that is robust and recovers experimental gene expression profiles. The Plant Cell 16:2923–2939. doi: https://doi.org/10.1105/tpc.104.021725

Gierer, A., and Meinhardt, H. (1972). A theory of biological pattern formation. Kybernetik 12:30–39. doi: https://doi.org/10.1007/BF00289234

Goto, K., and Meyerowitz, E.M. (1994). Function and regulation of the Arabidopsis floral homeotic gene PISTILLATA. Genes Dev 8:1548–1560. doi: 10.1101/gad.8.13.1548

Gruel, J., Landrein, B., Tarr, P., Schuster, C., et al. (2016). An epidermis-driven mechanism positions and scales stem cell niches in plants. Sci. Adv. 2: e1500989. doi: 10.1126/sciadv.1500989

Haspolat, E. Huard, B. and Angelova, M. (2019). Deterministic and Stochastic Models of Arabidopsis thaliana Flowering. Bulletin of Mathematical Biology 81:277–311. doi: https://doi.org/10.1007/s11538-018-0528-x

Hepworth, S.R., Klenz, J.E., and Haughn, G.W. (2005). UFO in the Arabidopsis inflorescence apex is required for floral-meristem identity and bract suppression. Planta 223:769–778. doi: 10.1007/s00425-005-0138-3

Ikeda, M., Mitsuda, N., and Ohme-Takagi, M. (2009). Arabidopsis WUSCHEL is a bifunctional transcription factor that acts as a repressor in stem cell regulation and as activator in floral patterning. The Plant Cell 21:3493–3505. doi: 10.1105/tpc.109.069997

Ingram, G.C. (2004). Between the sheets: inter-cell-layer communication in plant development. Phil. Trans. R. Soc. Lond. B 359: 891–906. doi: 10.1098/rstb.2003.1356

Jack, T. (2004). Molecular and Genetic Mechanisms of Floral Control. The Plant Cell 16:s1–s17. doi: https://doi.org/10.1105/tpc.017038

Jaeger, J., Surkova, S., Blagov, M., Janssens, H., Kosman, D., Kozlov, K.N., Myasnikova, M.E., Vanario-Alonso, C.E., Samsonova, M., Sharp, D.H., and Reinitz, J. (2004). Dynamic control of positional information in the early Drosophila embryo. Nature 430:368–371. doi: 10.1038/nature02678

Jönsson, H., Heisler, M., Reddy, G.V., Agrawar, V., Gor, V., Shapiro, B.E., Mjölsness, E., and Meyerowitz, E.M. (2005). Modeling the organization of the WUSCHEL expression domain in the shoot apical meristem. Bioinformatics 21:pSuppl 1, i232-i240. doi: 10.1093/bioinformatics/bti1036

Liu, X., Kim, Y.J., Müller, R., Yumu, R.E., Liu, C., Pan, Y., Cao, X., Goodrich, J., and Chen, X. (2011). AGAMOUS Terminates Floral Stem Cell Maintenance in Arabidopsis by Directly Repressing WUSCHEL through Recruitment of Polycomb Group Proteins. The Plant Cell 23:3654–3670. doi: 10.1105/tpc.111.091538

Maizel, A., and Weigel, D. (2004). Temporally and spatially controlled induction of gene expression in Arabidosis thaliana. The Plant Journal 38:164–171. doi: https://doi.org/10.1111/j.1365-313X.2004.02027.x

Mandel, M.A., Bowman, J.L., Kempin, S.A., Ma, H., Meyerowitz, E.M., and Yanofsky, M.F. (1992). Manipulation of flower structure in transgenic tobacoo. Cell 71:133–143. doi: https://doi.org/10.1007/BF00013745

Mendoza, L., Thieffry, D., and Alvarez-Buylla, E.R. (1999). Genetic control of flower morphogenesis in Arabidospsis thaliana: a logical analysis. Bioinformatics 15:593. doi: 10.1093/bioinformatics/15.7.593

Moyroud, E., Gómez-Minguet, E., Ott, F., Yant, L., Posé, D., Monniaux, M., Blanchet, S., Bastien, O., Thévenon, E., Weigel, D., Schmid, M., and Parcy, F. (2001). Prediction of regulatory interactions from genome sequences using a biophysical model for the Arabidopsis LEAFY transcription factor. The Plant Cell 23:1293–1306. doi: https://doi.org/10.1105/tpc.111.083329

Ng, M., and Yanofsky, M.F. (2001). Activation of the Arabidopsis B Class Homeotic Genes by APETALIA1. The Plant Cell 13:739–753. doi: 10.1105/tpc.13.4.739

Okamuro, J.K., Den Boer, B.G.W., Lotys-Prass, C., Szeto, W., and Jofuku, K.D. (1996). Flowers into shoots: Photo and hormonal control of a meristem identity switch in Arabidopsis. Proc. Natl. Acad. Sci. USA 93: 13831–13836. doi: https://doi.org/10.1073/pnas.93.24.13831

Parcy, F., Bomblies, K., and Weigel, D. (2002). Interaction of LEAFY, AGAMOUS and TERMINAL FLOWER1 in maintaining floral meristem identity in Arabidopsis. Development 129:2519–2527.

Pelaz, S., Ditta, G.S., Baumann, E., Wisman, E., and Yanofsky, M.F. (2000). B and C floral organ identity functions require SEPALLATA MADS box genes. Nature 405: 200–203. doi: 10.1038/35012103

Pelaz, S., Tapia-Lopez, R., Alvarez-Buylla, E.R., and Yanofsky, M.F. (2001). Conversion of leaves into petals in Arabidopsis. Curr. Biol. 11: 182–184. doi: 10.1016/s0960-9822(01)00024-0

Pidkowich, M.S., Kienz, J.E., and Haughn, G.W. (1999). The making of a flower: control of floral meristem identity in Arabidopsis. Trends in Plant Science 4:64–70. doi: 10.1016/s1360-1385(98)01369-7

Prigogine, I., and Nicolis, G. (1967). On symmetry-breaking instabilities in dissipative systems. J. Chem. Phys. 46:3542–3550. doi: 10.1063/1.1841255

Prigogine, I., and Lefever, R. (1968). Symmetry breaking instabilities in dissipative systems. II. J. Chem. Phys. 48:1695–1700. doi: https://doi.org/10.1063/1.1668896

Sablowsky, R. (2009). Cytokinin and WUSCHEL tie the knot around plant stem. PNAS 106:16016–16017.

Samach, A., Klenz, J.E., Kohalmi, S.E., Risseeuw, S.E., Haughn, G.W., and Crosby, W.L. (1999). The UNUSUAL FLORAL ORGANS gene of Arabidopsis thaliana is an F-box protein required for normal patterning and growth in the floral meristem. The Plant Journal 20:433–445. doi: 10.1046/j.1365-313x.1999.00617.x

Scheres, B. (1998). A LEAFY link from outer space. Nature 395:545–547. doi: https://doi.org/10.1038/26858

Sessions, A., and Yanofsky, M.F. (2000). Cell-Cell Signaling and Movement by the Floral Transcription Factors LEAFY and APETALA1. Science 289:779–781. doi: 10.1126/science.289.5480.779

Stewart, D., Graciet, E., and Wellmer, F. (2016). Molecular and regulatory mechanisms controlling floralorgan development. The FEBS Journal 283:1823–1830. doi: https://doi.org/10.1111/febs.13640

Traas, J., and Vernoux, T. (2002). The shoot apical meristem: the dynamics of stable structure. Phil. Trans. R. Soc. Lond. B 357:737–747. doi: 10.1098/rstb.2002.1091

Turing, A.M. (1952). The chemical basis of morphogenesis. Philosophical Transactions of the Royal Society of London. Series B, Biological Sciences 237:37–72. doi: https://doi.org/10.1098/rstb.1952.0012

Villarreal, C., Padilla-Longoria, P., and Alvarez-Buylla, E.R. (2012). General Theory of Genotype to Phenotype Mapping: Derivation of Epigenetic Landscapes from N-Node Complex Gene Regulatory Networks. Physical Reviews Letters 109:118102. doi: 10.1103/PhysRevLett.109.118102

Wang, C.C.N., Chuang, P., Ng, K., Chang, C., Sheu, P.C.Y., and Tsai, J.P. (2014). A model comparison study of the flowering time regulatory network in Arabidopsis. BMC Systems Biology 8:15. doi: https://doi.org/10.1371/journal.pcbi.1007671

Wolpert, L. (1994). Positional information and pattern formation in development. Developmental Genetics 15:485–490. doi: 10.1002/dvg.1020150607

Wu, X., Dinneny, J.R., Crawford, K.M., Rhee, Y., Citovsky, V., Zambryski, P.C., and Weigel, D. (2003). Modes of intercellular transcription factor movement in the Arabidopsis apex. Development 130:3735–3745. doi: 10.1242/dev.00577

